# CryoID2: deep learning-enhanced identification of protein and RNA from cryoEM maps

**DOI:** 10.64898/2026.11.03.764069

**Authors:** Qibo Xu, Xinye Yu, Leon Wu, Shi Feng, Z. Hong Zhou

## Abstract

Identifying the molecular components of cryoEM/cryoET maps of endogenous or poorly characterized complexes remains a major bottleneck. CryoID2 addresses this by integrating deep learning–based feature prediction with sequence prioritization and structural evaluation. Three 3D U-Nets extract structure classes, key-atom positions, and reduced residue/nucleotide types from the input map. These predictions support backbone tracing and the generation of short sequence queries that rank candidates from a user-supplied pool—using BLASTp for proteins and a maximin strategy for RNA. Corresponding PDB or AlphaFold models are then retrieved, rigidly fitted using predicted features and experimental density, and ranked by map agreement. Module-level tests show high-recall feature prediction and resolution-dependent atom clustering, while a 52-case benchmark recovers near-reference fitting quality. In practice, cryoID2 correctly identified PfA-M17 from a 166-protein mass-spectrometry pool and the unannotated Ca-raiA RNA from a genome-derived non-coding sequence set, demonstrating a unified sequence-plus-structure route to molecular identification in cryoEM maps.

## INTRODUCTION

The resolution revolution in single-particle cryogenic electron microscopy (cryoEM) has enabled structural determination of macromolecules that are difficult to study by conventional methods, including membrane proteins, conformationally heterogeneous assemblies, and complexes isolated directly from cells [1–3]. By imaging particles in a frozen-hydrated state and computationally resolving heterogeneous populations, cryoEM can preserve biologically relevant conformations while accommodating limited sample amounts and compositional heterogeneity [e.g., 4–9]. In parallel, cellular cryogenic electron microscopy (cryoET) has advanced to deliver near-atomic-resolution structures of macromolecular complexes directly inside cells, enabling molecular sociology of native cellular assemblies [see reviews 10–12].

These advantages have broadened the range of native macromolecular assemblies accessible to structural analysis. However, reconstructing a three-dimensional density map does not by itself establish the molecular identity of the component represented by the map.

This problem is particularly challenging for maps reconstructed from endogenous or incompletely characterized samples such as crude cellular extracts and intact (or lamellae of) cells. Proteomic or transcriptomic analyses may provide hundreds to thousands of plausible protein or RNA candidates, yet they do not establish which candidate corresponds to a particular three-dimensional density map. At near-atomic resolution, well-resolved local features such as amino-acid side chains can support sequence assignment. At lower or spatially heterogeneous resolution, however, backbone and secondary-structure features may remain visible while residue-level identities become ambiguous [4]. Manual molecular identification therefore requires substantial expertise and can become a major bottleneck when many candidate molecules, unknown complexes, or conformational states must be evaluated.

The original cryoID method (*i.e.*, cryoID1 hereafter) addressed this identification problem within a bottom-up structural-proteomics workflow [4]. CryoID1 selected well-resolved regions of a cryoEM map, traced local protein backbones, assigned approximate residue identities, and reduced the 20 amino acids to a six-letter alphabet based on similarities in side-chain densities. Searching these error-tolerant sequence fragments against a mass spectrometry-derived candidate pool enabled protein identification without prior knowledge of the sequence represented by the map. This work demonstrated that density-derived sequence information could link an unknown cryoEM reconstruction to its molecular identity. However, the cryoID1 workflow was developed primarily for protein identification and relied mainly on algorithmic feature extraction and sequence-based matching, leaving opportunities to incorporate deep learning-based feature prediction, extend the approach to RNA, and integrate structural evaluation into the identification process.

Subsequent deep-learning methods have further advanced automated interpretation of cryoEM maps. DeepTracer predicts backbone and residue-level features for de novo protein modeling [13]. ModelAngelo performs molecular identification through profile hidden Markov model searches based on predicted sequence information [14]. DeepMainmast combines learned atom and residue predictions, backbone tracing, and AlphaFold2 models for protein structure modeling [15]. Cryo2Struct combines transformer-based atom and residue prediction with HMM-based sequence alignment to construct protein backbones [16]. CryoAtom adapts AlphaFold2-inspired attention mechanisms to improve protein model building from density maps [17]. For RNA, EMRNA combines nucleotide detection, backbone tracing, and sequence and secondary-structure information to construct all-atom models [18]. These approaches demonstrate that learned representations can automate important steps in atomic model interpretation and reduce manual intervention. However, de novo model building and molecular identification address related but distinct questions. Whereas de novo methods aim primarily to construct atomic models from cryoEM density, cryoID-based approaches focus on determining which candidate molecule is represented by an unknown cryoEM map.

The growth of public structure databases also provides an opportunity to incorporate structural information into molecular identification. The Protein Data Bank provides experimentally determined structures, while AlphaFold has greatly extended structural coverage for proteins lacking deposited experimental models [19–21]. Experimentally determined or computationally predicted RNA structures can likewise provide candidate models for structure-based evaluation when available [22]. However, selecting an appropriate structural model and placing it into an unknown cryoEM map remains nontrivial. Even sequence-identical PDB entries may represent different conformations, ligand-bound states, constructs, or assemblies. Candidate models therefore must be evaluated against experimental density and other map-derived structural features, allowing structural agreement to further discriminate among sequence-prioritized candidates during molecular identification.

Here, we present cryoID2, a deep learning-enhanced method that extends the cryoID1 framework to protein and RNA identification through integrated sequence and structural analysis. Starting from an unknown cryoEM map and a user-defined candidate sequence pool, cryoID2 uses three-dimensional neural networks to predict voxel-wise structural and sequence-informative features, including structure classes, key-atom positions, and reduced residue or nucleotide types. These predicted features are organized through backbone tracing and graph-based procedures to generate short sequence queries, which are then used to prioritize candidate sequences using mode-specific ranking strategies. Protein candidates are ranked using BLASTp-based scoring [23], whereas RNA candidates are prioritized using an RNA-specific sequence-ranking strategy. Experimental or computationally predicted structural models corresponding to prioritized candidates are then retrieved or provided, fitted to the cryoEM map using the predicted structural features, and ranked according to their agreement with the map, enabling structural information to further refine candidate selection.

By integrating deep learning-based feature prediction with sequence-based candidate prioritization and structure-based evaluation, cryoID2 extends the cryoID1 strategy from protein-focused sequence identification to an integrated protein and RNA identification framework.

Rather than replacing de novo model-building methods, cryoID2 addresses the related but distinct problem of molecular identification by leveraging both map-derived sequence information and the agreement of candidate structural models with experimental cryoEM density. This integration reduces manual transitions between map interpretation, candidate sequence analysis, and structural evaluation, providing a more systematic approach for identifying unknown molecular components in cryoEM maps.

## RESULTS

### cryoID2 integrates sequence and structural evidence for protein and RNA identification

cryoID2 extends the cryoID1 framework [4] into an integrated identification workflow that combines deep learning-based feature prediction, sequence-based candidate prioritization, and structure-based evaluation. The pipeline takes two primary inputs: an unidentified cryoEM density map and a user-defined candidate sequence pool, which may include proteins identified by mass spectrometry or RNA candidates derived from transcriptomic or genomic information. Protein and RNA candidates are processed using distinct sequence-ranking strategies, after which structural models corresponding to prioritized candidates are fitted to the experimental cryoEM density and ranked according to their map agreement, further refining candidate selection.

Overall, cryoID2 consists of four major steps: deep learning-based feature prediction, query generation, mode-specific candidate ranking, and structural model evaluation (Fig. 1). First, neural network-based prediction is applied to the input cryoEM map to identify local structural and sequence-informative features, including structure classes, key-atom positions, and reduced residue or nucleotide types. Second, predicted key atoms and structural features are used to reconstruct backbone segments and generate short sequence queries. Third, protein and RNA candidates are evaluated using distinct sequence-based ranking strategies tailored to their respective sequence characteristics. Finally, structural models corresponding to selected candidates are retrieved or provided by the user, fitted to the cryoEM map using the predicted structural features and experimental density, and ranked according to their map agreement.

**Figure 1.**
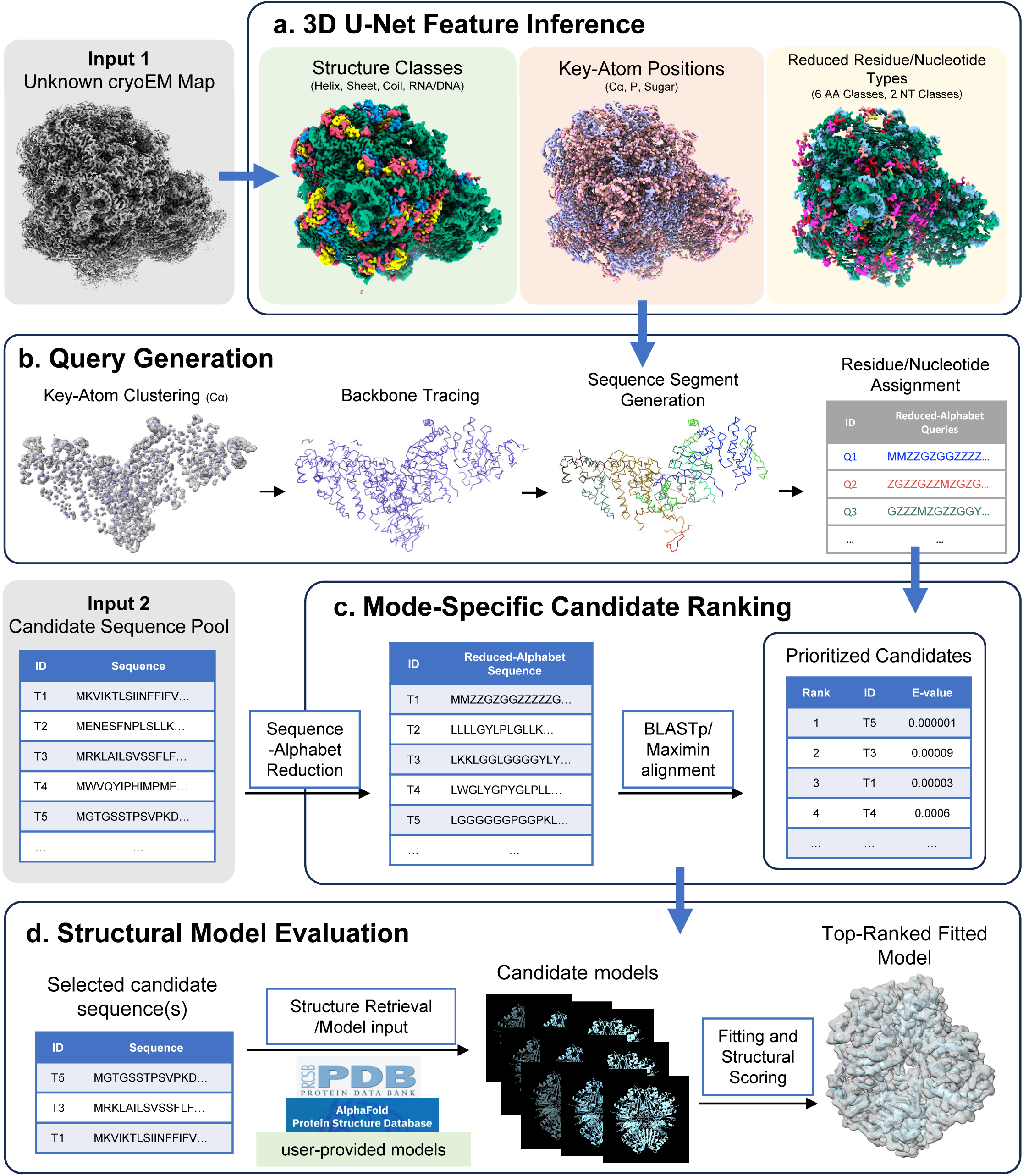
Overview of the cryoID2 workflow. CryoID2 takes an unknown cryoEM map and a user-defined candidate sequence pool as inputs. (a) U-Net-based neural networks predict structural and sequence-informative features, including structure classes, key atom positions, and reduced residue or nucleotide types. (b) Predicted key atoms are clustered and traced to reconstruct backbone segments, along which reduced-alphabet residue or nucleotide types are assigned to generate short sequence queries. (c) Candidate sequences are recoded into the corresponding reduced alphabet and prioritized using mode-specific sequence-ranking strategies, including BLASTp-based ranking for proteins and maximin ranking for RNA. (d) Structural models corresponding to selected candidates are retrieved from structure databases or provided by the user, fitted to the cryoEM map using predicted structural features and experimental density, and ranked according to their structural agreement with the map.

The graphical interface exposes the same workflow through project creation, density-map and candidate-pool selection, run controls, progress reporting, and execution logs (Fig. 2a). The sequence ranking result opens the ranked candidates and their sequence scores (Fig. 2b), and the model result opens fitted protein or RNA models with their density in UCSF ChimeraX [24] (Fig. 2c). The GUI translates user selections into command-line arguments and invokes the same cryoID2 pipeline, providing a graphical front end to the command-line interface.

**Figure 2.**
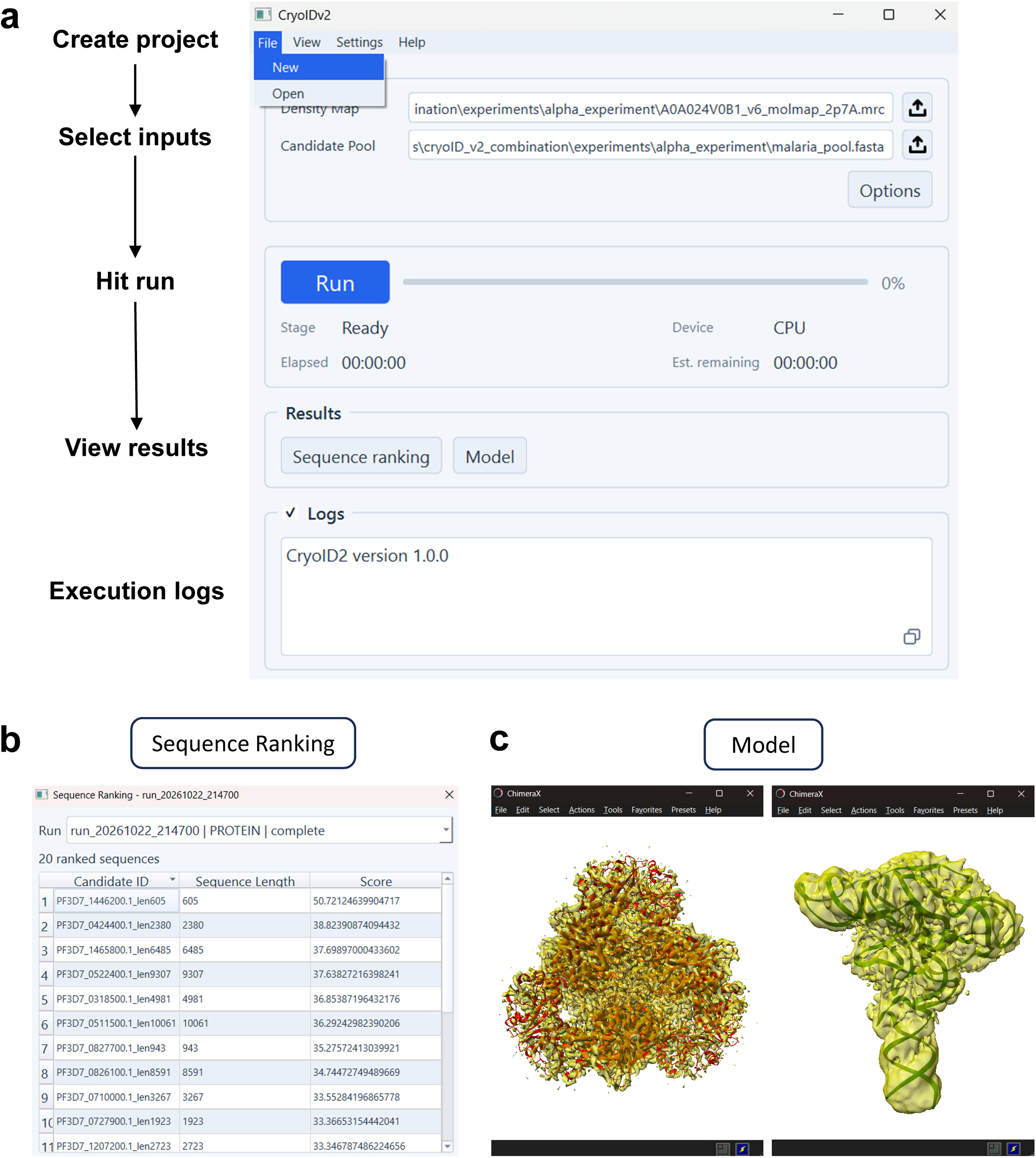
CryoID2 graphical user interface and output visualization. The cryoID2 GUI provides a user-friendly interface for sequence identification from cryoEM density maps. (a) Overview of the CryoID2 workflow. (b) Ranked candidate protein sequences with matching scores after clicking the sequence ranking button. (c) Visualization of the resulting protein and RNA models in ChimeraX after clicking the model button.

### Deep learning-based feature prediction

The first step of cryoID2 is deep learning-based feature prediction from the input cryoEM density map (Fig. 1a). The unidentified map is analyzed using a set of U-Net-based neural networks [25,26] that predict multiple structural and sequence-informative features at the voxel level.

These predicted features include structure classes, key-atom positions, and reduced residue or nucleotide types. Here, reduced residue or nucleotide types refer to simplified sequence alphabets in which chemically or structurally similar residues or nucleotides are grouped into a smaller number of classes. For proteins, the 20 standard amino acids are represented by six reduced amino acid classes, whereas nucleotides are represented by two reduced nucleotide classes. This reduced representation simplifies voxel-level classification from cryoEM density maps and makes the generated queries more tolerant of local prediction uncertainty. These voxel-level predictions provide the foundation for subsequent backbone tracing, residue or nucleotide assignment, and sequence query generation.

### Query generation

After feature prediction, cryoID2 uses the predicted key-atom positions to reconstruct backbone traces. For proteins, predicted Cα positions are obtained by clustering U-Net Cα-probability voxels into discrete positions and are subsequently connected to form backbone segments. For RNA, predicted phosphate and sugar key-atom positions are similarly used to trace nucleic-acid backbone segments. These traced segments define the spatial order of residues or nucleotides in the cryoEM map. Because positional information alone can permit multiple geometrically plausible connections between neighboring key atoms, tracing is further guided by U-Net-predicted structural regions that serve as a soft structural scaffold. Helix predictions are used to guide protein backbone tracing, whereas the predicted RNA/DNA structural class is used for RNA backbone tracing.

Reduced-alphabet residue or nucleotide types are then assigned along the traced backbone segments to generate short sequence queries (Fig. 1b) that capture local sequence information predicted from the cryoEM map. The reduced alphabet decreases the impact of local prediction uncertainty and density ambiguity while preserving sequence-informative signals for downstream candidate ranking.

### Mode-specific candidate ranking

Protein and RNA candidate pools are ranked using distinct sequence-based strategies tailored to their respective sequence characteristics (Fig. 1c). For protein candidate ranking, map-derived query fragments and candidate sequences are recoded into the same reduced amino acid alphabet and compared using BLASTp-short [23]. Evidence from multiple query fragments is integrated into a composite E-value for each candidate, allowing candidates to be ranked according to their overall agreement with the complete set of map-derived queries. Protein candidates with lower composite E-values, reflecting stronger combined sequence-match evidence across the query set, receive higher priority for downstream structural evaluation.

For RNA candidate ranking, map-derived query fragments and candidate sequences are recoded into the same two-letter reduced nucleotide alphabet. Because RNA’s two-letter reduced alphabet provides less sequence discrimination than the six-letter protein alphabet, we used a different method to favor candidates with consistent rather than combined support across query fragments. The method is termed maximin because it prioritizes candidates that maximize the minimum query-specific percentile score across all query fragments. Each query is compared with every candidate using ungapped sliding-window matching in both forward and reversed orientations, and the highest local sequence identity is retained for each query-candidate pair. For each query, the retained identity values are converted to percentile ranks across the candidate pool, with tied values assigned to their average rank. Each candidate’s maximin score is its lowest percentile across all queries, and candidates are ranked in descending order of this score. Thus, a strong match to one query cannot compensate for weak relative support from another. Ties are resolved by higher mean percentile followed by higher mean sequence identity.

For both protein and RNA analyses, a configurable number or percentile of the highest-ranking candidates is passed to the subsequent structural evaluation stage.

### Structural model evaluation

Following sequence-based candidate prioritization, cryoID2 retrieves or accepts structural models corresponding to selected candidates for structure-based evaluation against the cryoEM map (Fig. 1d). For proteins, selected sequences are matched against PDB protein sequences to identify corresponding experimental structures, and available UniProt mappings are used to retrieve AlphaFold Database models [21,27]. For RNA, selected sequences are searched against a PDB-derived RNA sequence database to identify available experimental structures.

Each candidate structural model is then fitted to the cryoEM map using deterministic rigid-body optimization guided by U-Net-predicted key-atom and secondary-structure fields together with the experimental map density. Initial poses are generated from high-confidence predicted structural features and subsequently refined to maximize a composite whole map and predicted feature agreement score. Fitted models are ranked according to this score, and the highest-scoring models are retained for structural comparison. Thus, sequence-based ranking first narrows the candidate pool, while map-based structural evaluation further discriminates among the selected candidates.

### Benchmarking evaluates the performance of key cryoID2 modules

To systematically evaluate cryoID2, we benchmarked the performance of its major computational modules rather than assessing the method solely on the final molecular identification outcome. Because cryoID2 consists of multiple sequential steps (Fig. 1), evaluating individual modules separately allows us to characterize the performance of feature prediction, query generation, candidate prioritization, and structural evaluation.

We first evaluated the deep learning module for voxel-level prediction of structural and sequence-informative features from cryoEM density maps (Fig. 3a). Three U-Net models (U-Net 1, 2, and 3) were assessed, corresponding respectively to the prediction of structure classes, key-atom positions, and reduced residue or nucleotide types. The structure-class model achieved a voxel-wise accuracy of 0.86, with precision, recall, and F1 score of 0.24, 0.97, and 0.38, respectively. The key-atom model achieved the highest voxel-wise accuracy of 0.95, with precision, recall, and F1 score of 0.16, 0.99, and 0.27, respectively. The reduced residue or nucleotide types model achieved a voxel-wise accuracy of 0.86, with precision, recall, and F1 score of 0.22, 0.92, and 0.36, respectively. Across all three models, voxel-wise recall was consistently high, whereas precision was lower, indicating sensitive detection of relevant map features together with additional voxel-level predictions that are subsequently refined through clustering, tracing, and downstream candidate-ranking procedures.

**Figure 3.**
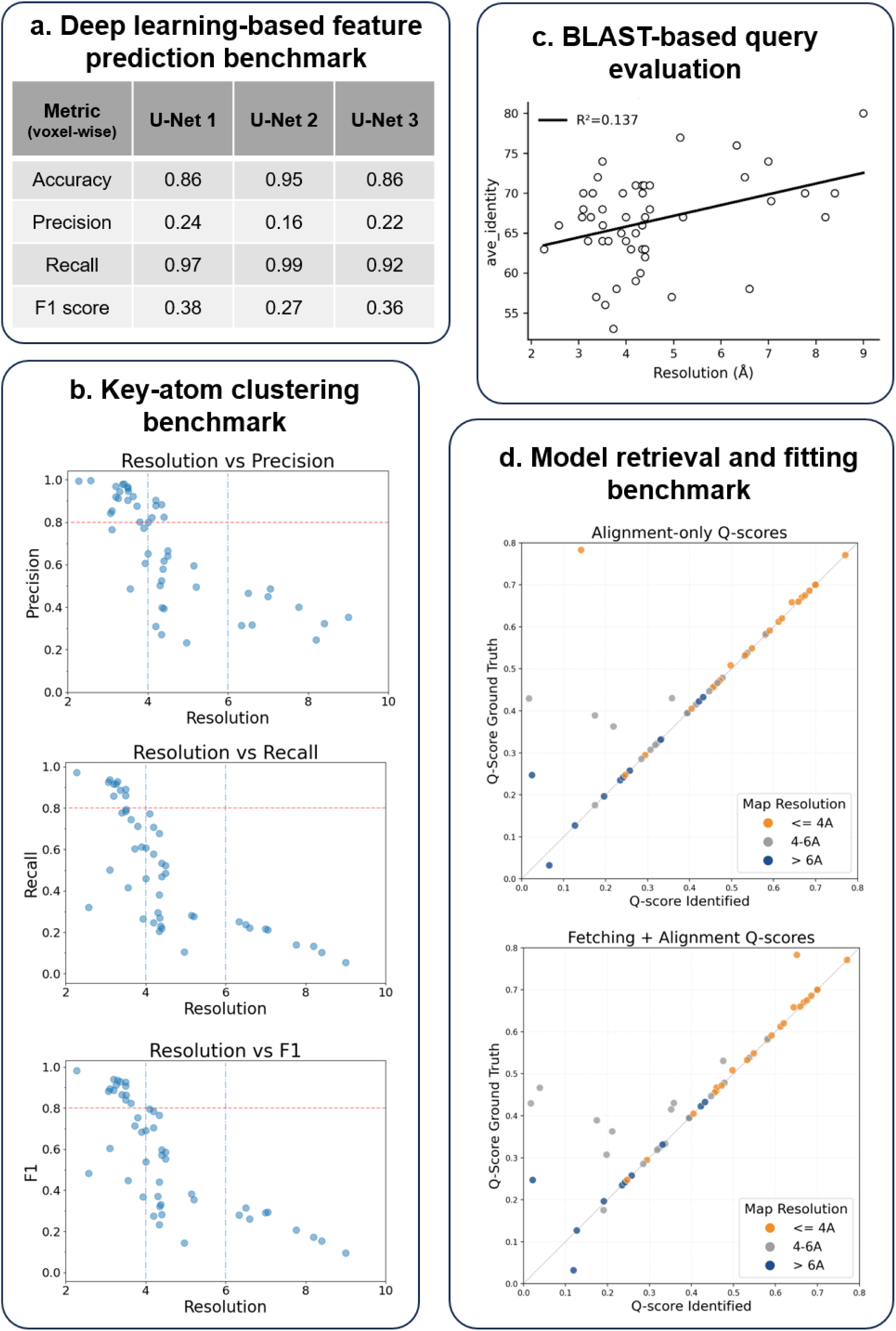
Benchmarking of key cryoID2 modules. (a) Voxel-wise performance of three U-Net models used for deep learning-based feature prediction from cryoEM density maps. The evaluated models predict structure classes, key atom positions, and six-class reduced residue types, respectively. Metrics include accuracy, precision, recall, and F1 score. (b) Key atom clustering benchmark showing precision, recall, and F1 score as functions of map resolution. The clustering step converts voxel-level key atom predictions into discrete atom candidates for downstream backbone tracing and query generation. (c) BLASTp-based query evaluation showing the relationship between map resolution and average sequence identity of generated queries. (d) Structural model retrieval and fitting benchmark comparing Q-scores of cryoID2-identified models with deposited ground-truth models across maps with different resolution ranges.

We next evaluated the key-atom clustering module, which converts voxel-level predictions into discrete atom candidates for backbone tracing and query generation (Fig. 3b). Because accurate Cα localization is essential for protein backbone reconstruction, clustering performance was assessed using precision, recall, and F1 score across maps spanning a range of resolutions. All three metrics showed a clear resolution-dependent trend, with stronger clustering performance for higher-resolution maps and progressively lower performance for lower-resolution maps. These results demonstrate that map resolution influences both recovery of true Cα positions and the suppression of false-positive atom candidates, and that voxel-level prediction performance does not directly translate into atom-level reconstruction accuracy.

We then evaluated the quality of sequence queries generated from the reconstructed backbone (Fig. 3c). Because unrecovered Cα positions introduced gaps in the reconstructed backbone, each protein sample yielded one or more sequence queries, which were searched against the corresponding reference sequence using BLASTp to assess sequence similarity. The reference sequence was obtained from the corresponding atomic model. The average sequence identity of the generated queries showed only a weak dependence on map resolution, with an R² of 0.137. This indicates only a weak relationship between query sequence identity and cryoEM map resolution.

Finally, we evaluated structural model retrieval and map-based evaluation using a curated set of 52 random published protein cryoEM cases (Fig. 3d; Methods). To limit redundancy, the benchmark included only one case per randomly selected gene. In the fetch-plus-alignment benchmark, each input protein sequence was matched against PDB chain sequences to retrieve corresponding PDB structures and chain-associated AlphaFold Database models. All candidate models were then displaced from their initial poses before fitting and scoring. The selected PDB identifier exactly matched the deposited model in 40 cases. In the remaining 12 cases, the selected structure had a different PDB identifier but the same protein sequence as the reference model. The gene name, exact reference and selected PDB identifiers, together with the Q-scores for each case [28], are reported in Supplementary Table 1.

Across the 52 cases, the mean Q-score decreased from 0.4466 for the deposited models to 0.4120 for the cryoID2-selected models after fitting, corresponding to a mean difference of −0.0346. The median difference was −0.0001, indicating that the average decrease was dominated by a small number of failure cases with large Q-score losses.

To distinguish the contribution alignment and evaluation from upstream candidate retrieval, we performed a paired alignment-only control in which the deposited ground-truth model itself was randomly displaced and realigned using the same procedure. The mean Q-score after re-alignment was 0.4138, corresponding to a mean difference of −0.0328 relative to the deposited models, with the near-zero median difference of −0.0001. Together, the near-zero median differences and lower mean Q-scores indicate that fitting generally preserved reference-level map-model agreement, with substantial losses concentrated in a subset of cases. The gene name and exact Q-scores for each case are reported in Supplementary Table 2. Similar decreases in the two benchmarks suggest that the shared alignment and evaluation procedure accounts for much of the average loss, with candidate retrieval contributing relatively little additional loss.

### cryoID2 identifies an unknown protein from a mass spectrometry-derived candidate pool

After benchmarking the individual computational modules, we evaluated an end-to-end protein-identification case using the *P. falciparum* E1_D3 experimental map, treated as unidentified during analysis (Fig. 4a). A candidate pool was constructed from proteins identified by mass spectrometry, and cryoID2 was used to determine which candidate best matched the experimental map.

**Figure. 4.**
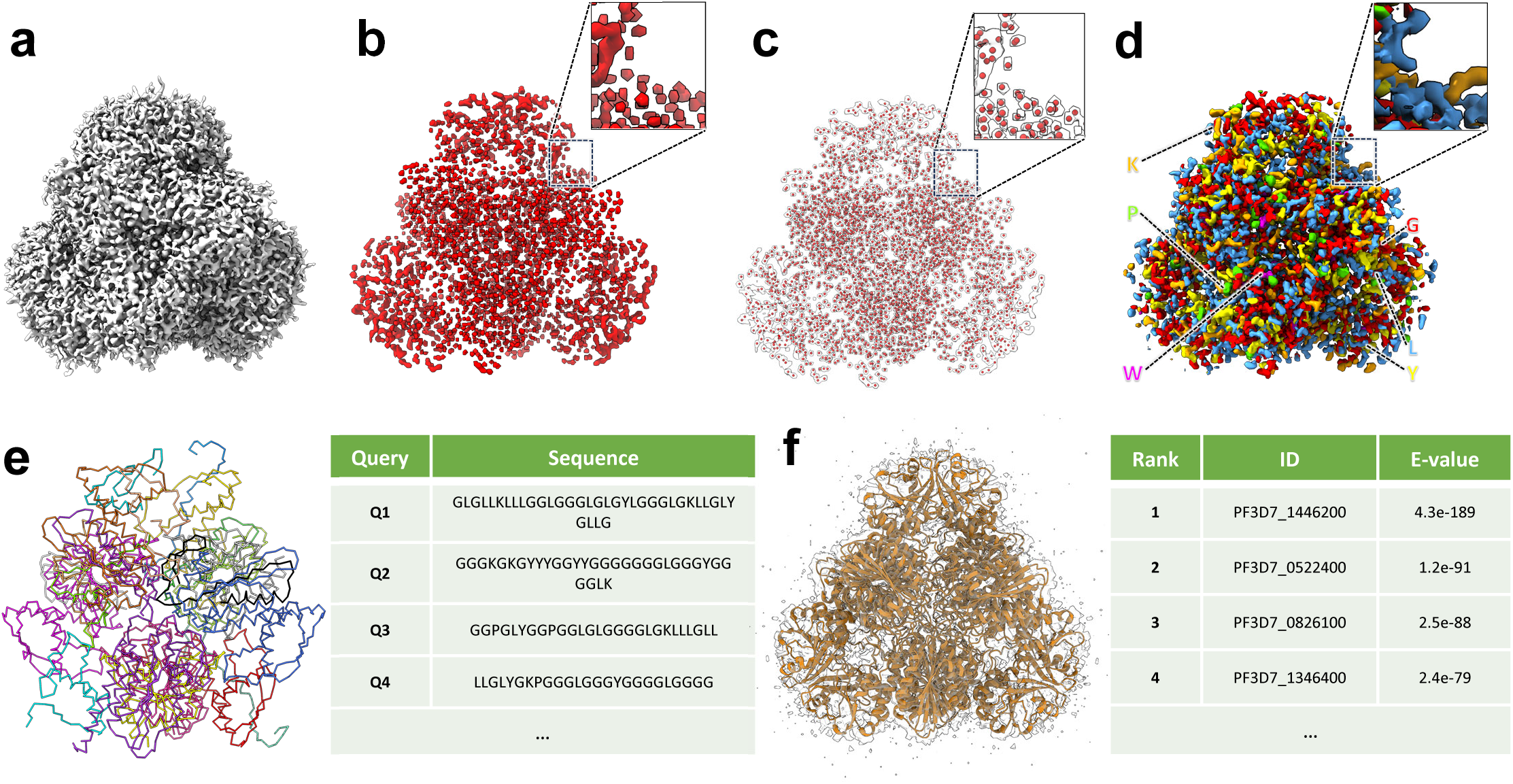
Protein identification from a mass spectrometry-derived candidate pool. (a) CryoEM map of the unidentified P. falciparum protein density. (b) CryoID2-predicted Cα density. (c) Clustered Cα atom positions with a zoom-in view. (d) Predicted six-class reduced residue types. (e) Map-derived sequence queries generated from backbone tracing and reduced residue-type assignment. (f) BLASTp-based candidate ranking identified PF3D7_1446200/PfA-M17, model PDB 4ZX8, as the top-ranked candidate.

The mass spectrometry-derived candidate pool contained 166 protein sequences. Starting from the *P. falciparum* cryoEM map, cryoID2 predicted Cα positions (Fig. 4b), clustered Cα positions into discrete atoms (Fig. 4c) and predicted reduced residue-type information (Fig. 4d), which were used for backbone tracing, sequence-query generation and alignment. In total, 20 map-derived sequence queries with an average length of 22.6 residues were generated, with a minimum sequence length parameter of 18 (Fig. 4e). The queries were generated in the six-class reduced amino acid alphabet, and the candidate protein sequences were recoded into the same alphabet for BLASTp-based ranking.

Sequence-based ranking placed PF3D7_1446200 first among the 166 mass spectrometry-derived candidates. This sequence corresponds to PfA-M17 and showed the strongest overall agreement with the map-derived queries. Structural models corresponding to this top-ranked candidate were then retrieved and evaluated against the cryoEM map. Among the retrieved models, a corresponding experimental PfA-M17 structure, PDB 4ZX8, achieved the highest structural score and was selected as the best-matching model at the recommended contour level of 0.25 (Fig. 4f).

Additionally, we compared the alignment of the top model from the top sequence, 4ZX8 (Supplementary Fig. 1a), to the second and third best ranked model from the top sequence and the top ranked models of the second and third ranked sequences. These models were 4ZY1 (Supplementary Fig. 1b), 4ZX9 (Supplementary Fig. 1c), 5LP8 (Supplementary Fig. 1d), and 9YGJ (Supplementary Fig. 1e), respectively [9,29,30]. The second and third ranked models for the top sequence were a much better fit than the best models for the second and third ranked sequences, providing additional structural support for the top sequence assignment.

After establishing the protein identity as PfA-M17, we performed two synthetic-density controls (Fig. 5a). Density was simulated at 2.7 Å from 4ZX8 chain A and AlphaFold Database model A0A024V0B1. CryoID2 retrieved A0A024V0B1 as the highest-ranked model for both the experimental-structure-derived density (Fig. 5b) and the AlphaFold-derived density (Fig. 5c).

**Figure. 5.**
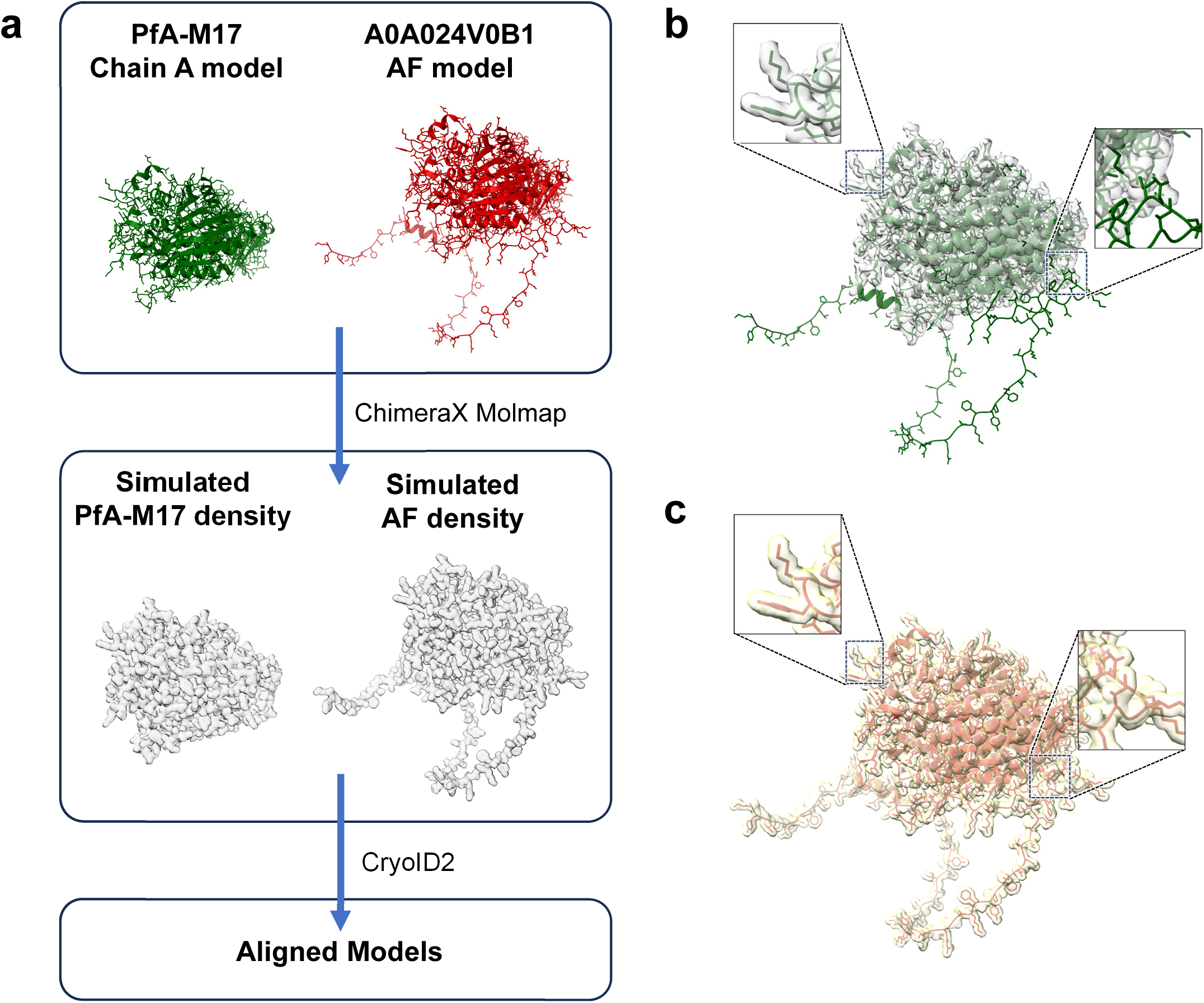
AlphaFold model retrieval and fitting in synthetic-density controls. (a) 2.7 Å density maps are generated from PfA-M17 chain A and A0A024V0B1 AlphaFold model, followed by model retrieval and fitting with cryoID2. (b) A0A024V0B1 is identified as the public best fit model and fitted to the experimental structure derived density. (c) A0A024V0B1 is identified as the public best fit model and fitted to AlphaFold structure derived density.

Regions of the AlphaFold model supported by the experimental-structure-derived density showed good agreement with the map (Fig. 5b). These controls demonstrate automated retrieval and alignment of AlphaFold Database models, supporting their use when suitable experimental models are unavailable.

This practical case demonstrates that cryoID2 can identify a specific protein from an experimentally constrained candidate pool and subsequently evaluate structural models associated with the identified protein. Together, these results illustrate the integration of map-derived sequence information with downstream structural evaluation in the cryoID2 protein-identification workflow.

### A proof-of-concept RNA case combines sequence-based prioritization with structural evaluation

We next evaluated RNA candidate identification using the 3.0 Å cryoEM map EMD-47655 from *Clostridium acetobutylicum* ATCC 824 (Fig. 6a), with the 204-nucleotide RNA modeled in PDB 9E73 serving as the reference target [31].

**Figure. 6.**
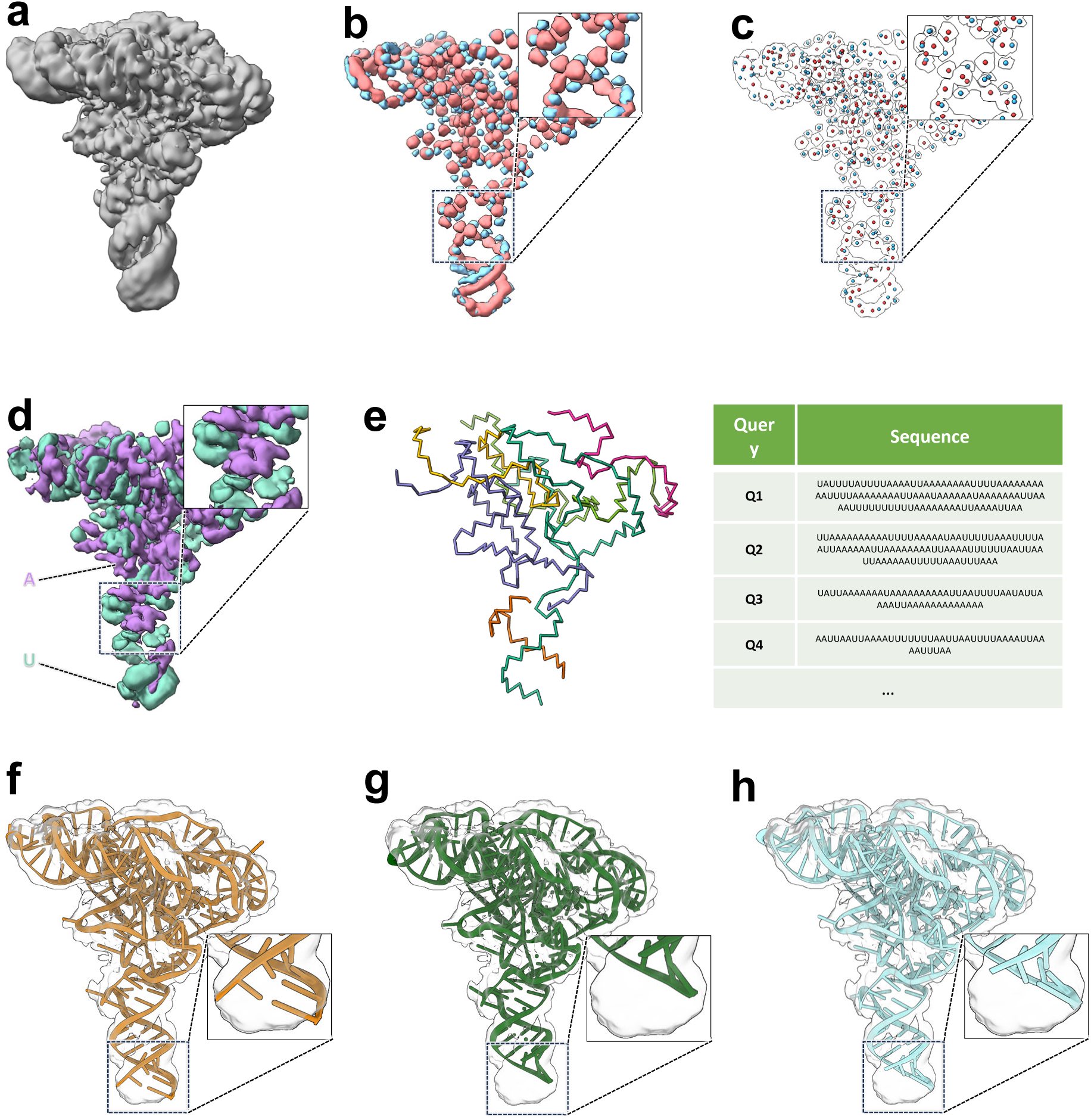
RNA identification from a genome-derived non-CDS candidate pool. (a) CryoEM map of EMD-47655. (b) CryoID2-predicted phosphate (blue) and sugar (red) density. (c) Clustered phosphate (blue) and sugar (red) atom positions with a zoom-in view. (d) Predicted two-class reduced nucleotide types. (e) Map-derived RNA query fragments generated from backbone tracing and reduced nucleotide-type assignment. (f) The deposited Ca-raiA RNA structure, PDB 9E73, was retrieved and selected as the best-matching structural model after fitting and structural evaluation. (g) The rank 2 model, PDB 9CXF. (h) The rank 3 model, PDB 9ELY.

We constructed the candidate pool directly from the complete *C. acetobutylicum* ATCC 824 genome assembly GCF_000008765.1, including chromosome NC_003030.1 and plasmid NC_001988.2 [32]. Because the target was known to be a non-coding RNA, annotated coding sequence (CDS) regions were excluded using the corresponding genome annotation. The remaining non-CDS genomic intervals of at least 50 nucleotides were retained. Each interval was represented by both its forward sequence and reverse complement, with thymine converted to uracil. This procedure yielded 4,632 candidate sequence records corresponding to 2,316 genomic intervals.

For this proof-of-concept analysis, the candidate pool was further constrained using the known 204-nucleotide reference length, retaining sequences within ±50% of this length. This produced a final search pool of 2,538 sequence records representing 1,269 genomic intervals. The target Ca-raiA sequence was contained within the reverse-complement representation of a 294-nucleotide chromosomal interval, together with flanking genomic sequence.

CryoID2 predicted phosphate and sugar density from the cryoEM map (Fig. 6b), clustered these predictions into discrete atom positions (Fig. 6c), assigned reduced nucleotide types along the resulting backbone traces (Fig. 6d), and generated six map-derived RNA query fragments (Fig. 6e). The query fragments and candidate sequences were represented using the same two-class reduced nucleotide alphabet. Using the RNA-specific maximin ranking procedure, the Ca-raiA-containing genomic interval ranked fourth in the sequence-based candidate ranking.

For structural evaluation, the top 50% of sequence-ranked candidates were searched against a PDB RNA-chain sequence database using BLASTN, yielding 15 nonredundant RNA models that were successfully fitted to the map. The ground truth Ca-raiA model, 9E73, ranked first (Fig. 6f), followed by 9CXF chain A (Fig. 6g) and 9ELY chain A (Fig. 6h) [33,34]. Despite their high structural similarity, cryoID2 assigned the deposited reference model a higher fitting score than the second- and third-ranked models, demonstrating discrimination among closely related RNA structures in this case.

Although the Ca-raiA-containing interval ranked fourth at the sequence-ranking stage, subsequent structure retrieval and structural evaluation further refined the candidate set and selected the deposited Ca-raiA structure as the highest-scoring retrieved model, illustrating how structural information can complement sequence-guided candidate prioritization in the cryoID2 RNA workflow.

## DISCUSSION

CryoID2 builds on the cryoID1 strategy for identifying molecular components directly from cryoEM maps. The original method demonstrated that approximate sequence information extracted from density could identify proteins within experimentally constrained candidate pools [4]. CryoID2 retains this principle while introducing deep learning-based feature prediction, extending the analysis to RNA, and incorporating structural comparison into the identification workflow. Map-derived sequence queries provide an initial basis for candidate prioritization, while available experimental or predicted models allow selected candidates to be examined against the experimental density. These developments extend cryoID1 beyond sequence matching without changing its central objective of determining which molecule is represented by an unidentified cryoEM map.

The modular benchmarks reveal different performance characteristics across the computational stages. The three U-Net models showed high voxel-level recall but substantially lower precision, emphasizing the importance of downstream processing when converting predicted features into discrete backbone traces and sequence queries. Cα clustering performance was associated with cryoEM map resolution, whereas query sequence identity showed only a weak linear relationship with resolution. Thus, the reported map resolution alone was not a strong linear predictor of query sequence identity in this analysis. In the separate 52-case structural benchmark, the median Q-score difference relative to deposited reference models was close to zero, although the mean difference was negative. Similar decreases in the two benchmarks suggest that the shared alignment and evaluation procedure account for much of the average loss, with candidate retrieval contributing relatively little additional loss. These results help distinguish the behavior of individual modules, but the structural benchmark should not be interpreted as an evaluation of end-to-end molecular identification across all 52 cases.

The contribution of structural information in cryoID2 extends beyond checking an identity already proposed by sequence analysis. This workflow complements several automated model-building approaches. DeepTracer predicts backbone atoms and residue types for protein modeling [13]. DeepMainmast integrates backbone tracing with AlphaFold2 models [15].

Cryo2Struct combines transformer-based atom and residue prediction with HMM-based sequence alignment [16], and CryoAtom adapts AlphaFold2-inspired attention mechanisms for protein model building [17]. For RNA, EMRNA integrates nucleotide detection, backbone tracing, and sequence and secondary-structure information to construct all-atom models [18]. ModelAngelo combines density, sequence, and local geometry for atomic-model construction and also identifies proteins of unknown sequence through profile hidden Markov model searches [14]. In cryoID2, sequence analysis first prioritizes candidates, after which corresponding structural models are retrieved, fitted to the map, and ranked using a composite score measuring agreement with predicted features and experimental density. Structural agreement therefore contributes to selecting among the retrieved candidate models beyond sequence matching. Reusing existing models also allows researchers to assess compatibility with previously determined structures and provides fitted models as starting points for further building and refinement. These methodological and practical differences do not by themselves establish superior identification performance. Direct comparisons under matched candidate-pool and map conditions would be needed to assess the relative strengths of these approaches.

The *P. falciparum* case illustrates how these stages operate in a practical protein-identification workflow. Map-derived queries placed PF3D7_1446200/PfA-M17 first among 166 mass spectrometry-derived candidates, after which models associated with this protein were retrieved and evaluated against the cryoEM map. The experimental PfA-M17 structure 4ZX8 [29] obtained the highest structural score. In this example, sequence ranking establishes the protein identity, whereas the structural identification stage selected among models associated with that identity. It did not independently compare all 166 protein identities. The AlphaFold synthetic-density tests were performed after identification and provided controlled checks of predicted-model retrieval and evaluation. They should therefore be distinguished from the initial identification experiment rather than treated as additional blind protein-identification cases.

Together, these analyses demonstrate the integration of sequence-based identification with downstream structural assessment while clarifying the contribution of each stage.

The RNA analysis provides an initial proof of concept for extending the cryoID strategy beyond proteins. In the Ca-raiA example, map-derived RNA queries ranked the Ca-raiA-containing genomic interval fourth within a genome-derived non-CDS candidate pool. BLASTN searches of the top 50% of sequence-ranked candidates against a PDB RNA-chain sequence database yielded 15 nonredundant RNA models that were successfully fitted to the cryoEM map. The deposited Ca-raiA structure, PDB 9E73 [31], received the highest structural score, followed by 9CXF chain A and 9ELY chain A. Despite the high structural similarity among these models, cryoID2 distinguished the deposited reference structure from the other leading candidates.

These results illustrate how structural retrieval and map-based evaluation can complement map-derived sequence ranking by comparing the structural compatibility of retrieved models. However, this analysis remains a proof of concept rather than a general benchmark of genome-scale RNA identification. The candidate pool was constrained using the known 204-nucleotide reference length, and only a single RNA case with an available deposited structure was evaluated. Performance may therefore depend on the definition of genomic intervals, prior constraints on candidate length, candidate-pool composition, and the availability of suitable structural models. Broader evaluation across RNA families, candidate-pool designs, and cryoEM map resolutions will be required to establish the general applicability of the approach.

The nucleic-acid branch provides a possible basis for future DNA identification because it uses an RNA/DNA structural class and a reduced alphabet that can accept thymine-containing input sequences. This software-level compatibility does not demonstrate that RNA-trained predictions generalize to DNA. DNA-specific feature prediction, tracing, candidate ranking, and structure-retrieval benchmarks would be needed before claiming DNA-identification performance.

A further direction is to extend cryoID2 toward automated interpretation of complex cryoEM maps using individual component models from public structural repositories and prediction databases. Combining image segmentation with map-derived feature predictions could help delineate density regions corresponding to individual molecular components, allowing identification and fitting to proceed within each region. Alignment of retrieved models could then make it practical to evaluate multiple candidate components and assemble their fitted structures into a model of the larger complex. Such a workflow could support the reuse of experimentally determined or predicted structures when a complete assembly model is unavailable. Realizing this approach would require methods for resolving overlapping density, accounting for repeated subunits and conformational differences, and evaluating both local fits and compatibility across the assembled complex. These extensions require dedicated validation beyond the identification examples presented here.

Several limitations remain relevant to the broader application of cryoID2. The reduced alphabet groups multiple residue or nucleotide identities into shared classes, trading sequence specificity for tolerance to uncertain predictions. Identification consequently depends on the information retained in the generated queries and on the composition of the candidate pool. Structural comparison is further limited by the availability and quality of candidate models. Its ranking applies to the models retrieved or supplied, not necessarily to every molecular identity in the original pool. In addition, the current rigid-body procedure adjusts model placement without accommodating internal conformational changes, which may limit agreement when a suitable sequence is represented by a different conformation. Future work could explore more flexible structural comparisons, sensitivity to candidate-pool composition, broader validation of RNA identification and extension to DNA, and automated modeling of complex assemblies through map segmentation, feature prediction, and rapid fitting of individual component models. Overall, cryoID2 extends the established cryoID approach by combining learned map features, sequence-based prioritization, and explicit structural comparison within a unified molecular-identification workflow.

## Supporting information

ALL Supplementary info

## RESOURCE AVAILABILITY

### Lead contact

Requests for further information and resources should be directed to and will be fulfilled by the lead contact, Z. Hong Zhou.

### Materials availability

This study did not generate new unique reagents.

### Data and code availability

cryoID2 is an open source program under the MIT license, available for download at https://github.com/t00shadow/cryoID_v2. The model weights are available at https://huggingface.co/cryoid2/cryoID2. Users do not need to manually download model weights; cryoID2 automatically downloads them on first run. See Key Resources Table for data availability.

## ACKNOWLEDGMENTS

This project is supported in part by National Institutes of Health through R01GM071940 (to ZHZ). We thank Michael Rebelo for discussion during the early stage of this project, Denver Harris for stress-testing cryoID2.

## AUTHOR CONTRIBUTIONS

Conceptualization (ZHZ, QX, XY), Data curation (QX, XY, LW), Formal analysis (QX, XY, LW, SF, ZHZ), Funding acquisition (ZHZ), Investigation (QX, XY, LW, SF, ZHZ), Methodology (QX, XY, LW, SF, ZHZ), Project administration (ZHZ, QX), Resources (ZHZ), Software (QX, XY, LW, SF, ZHZ), Supervision (QX, ZHZ), Validation (QX, XY, LW, SF), Visualization (QX, XY, LW, SF, ZHZ), Writing – original draft (QX, XY, LW, SF, ZHZ), Writing – review & editing (QX, XY, LW, ZHZ).

## DECLARATION OF INTERESTS

ZHZ is a cofounder of Nanomega cryoAI, a service company in Los Angeles, USA. Other authors declare no competing interest.

---

**Supplementary Figure 1. Ranked models fitted to the E1_D3 cryoEM density map.** (a) PDB 4ZX8, first-ranked model for the first-ranked protein sequence. (b) PDB 4ZY1, second-ranked model for the first-ranked protein sequence. (c) PDB 4ZX9, third-ranked model for the first-ranked protein sequence. (d) PDB 5LP8, first-ranked model for the second-ranked protein sequence. (e) PDB 9YGJ, first-ranked model for the third-ranked protein sequence.

**Supplementary Figure 2. Architecture and outputs of the three cryoID2 U-Net models.** (a) An input cryoEM density map is processed sequentially by three 19-layer 3D U-Net models. (b,c) U-Net 1 predicts voxel-wise structure classes, including helix, sheet, coil, RNA/DNA, and background. (d,e) U-Net 2 predicts key-atom positions, including protein Cα atoms, RNA phosphate atoms, and RNA sugar-ring atoms. (f,g) U-Net 3 predicts reduced residue or nucleotide types. Protein residues are represented by six reduced amino-acid classes, whereas RNA nucleotides are represented by two reduced nucleotide classes. The outputs of the preceding prediction stages are incorporated into the downstream U-Net inputs as indicated by the arrows.

**Supplementary Table 1. Structural retrieval and fitting performance in the 52-case fetch-plus-alignment benchmark.** Gene names, reference and retrieved PDB identifiers, and case-wise Q-scores are reported for deposited reference models and cryoID2-selected models. Retrieved candidate models were randomly displaced before fitting to the corresponding cryoEM maps. Q-scores measure map-model agreement following fitting and independent structural evaluation.

**Supplementary Table 2. Structural fitting performance in the 52-case alignment-only benchmark.** Gene names and case-wise Q-scores are reported for deposited reference models and the same models after random displacement and realignment to their corresponding cryoEM maps. Candidate retrieval was omitted, allowing changes in map-model agreement to be assessed for the shared alignment and evaluation procedure independently of retrieval.

**Supplementary Table 3. Metadata for the cryoID2 U-Net training dataset.** Metadata for the 730 cryoEM map–model pairs retained after quality-control filtering and used for cryoID2 dataset generation. Columns include EMDB entry title and identifier, map resolution, fitted PDB structure(s), UniProtKB and AlphaFold cross-references, Q-score, atom inclusion, recommended contour level, VOF, and Dice coefficient.

**Supplementary Table 4. Illustrative example of RNA candidate ranking.** Local sequence identities and corresponding percentile scores are shown for four hypothetical candidates compared against two queries. Each candidate’s final score is the lower of its two percentile scores. Higher final scores receive higher priority, favoring candidates supported consistently across queries. Candidate B ranks first because its lowest percentile, 0.75, is the highest among the four candidates.

## STAR★METHODS

### KEY RESOURCES TABLE

| REAGENT OR RESOURCE | SOURCE | IDENTIFIER |
|---|---|---|
| Deposited data |  |  |
| PfA-M17 experimental density map (E1_D3) | This paper | Pending |
| Ca-raiA cryoEM density map | EMDB | EMD-47655<br>https://www.ebi.ac.uk/emdb/EMD-47655 |
| *C. acetobutylicum* ATCC 824 genome and annotation | NCBI RefSeq | GCF_000008765.1; chromosome NC_003030.1; plasmid NC_001988.2<br>https://www.ncbi.nlm.nih.gov/datasets/genome/GCF_000008765.1/ |
| PDB sequence database for structure retrieval | PDB | pdb_seqres.txt.gz<br>https://files.rcsb.org/pub/pdb/derived_data/pdb_seqres.txt.gz |
| PfA-M17 experimental structure | PDB | PDB: 4ZX8<br>https://www.rcsb.org/structure/4ZX8 |
| Ca-raiA reference structure | PDB | PDB: 9E73; chain A<br>https://www.rcsb.org/structure/9E73 |
| PfA-M17 predicted structure | AlphaFold Protein Structure Database | AF-A0A024V0B1-F1-model_v6<br>https://alphafold.ebi.ac.uk/files/AF-A0A024V0B1-F1-model_v6.pdb |
| Pretrained cryoID2 U-Net model bundle | This paper | Structure-class, key-atom, and reduced residue/nucleotide models<br>https://huggingface.co/testaccount153/testupload/tree/3ae72b857d69e16dbfecc3af875733de55ef39a7/train_val_data |
| Software and algorithms |  |  |
| cryoID2 | This paper | https://github.com/t00shadow/cryoID_v2 |
| NCBI BLAST+ 2.17.0 | NCBI; Camacho et al. (2009) | https://ftp.ncbi.nlm.nih.gov/blast/executables/blast+/2.17.0/ |
| UCSF ChimeraX 1.9 | Pettersen et al. (2021) | https://www.cgl.ucsf.edu/chimerax/ |

## METHOD DETAILS

### Training data and voxel-label generation

Training data for the cryoID2 U-Net models were constructed using the cryoDataBot [35] dataset-generation pipeline. Following quality-control filtering, 730 cryoEM map-model pairs were retained for dataset construction (Supplementary Table 3). CryoEM maps were resampled to a uniform voxel size of 1 Å and paired with their corresponding deposited atomic models.

Ground-truth labels were generated from the atomic models on the same voxel grids as the processed cryoEM maps for the three prediction tasks used in cryoID2: structure classes, key-atom positions, and residue or nucleotide types. The structure classes comprised helix, sheet, coil, and RNA; the key-atom classes comprised protein Cα atoms, RNA phosphate atoms, and RNA sugar-ring atoms; and the residue or nucleotide labels initially comprised the 20 standard amino-acid types and four RNA nucleotide types.

For training the reduced residue or nucleotide type model, the original residue and nucleotide labels were subsequently recoded into the reduced alphabets used by cryoID2. The 20 amino acids were grouped into six classes: G/A/S/C/V/T/I, P, L/D/N/E/Q/M, K/R, H/F/Y, and W, whereas the four RNA nucleotides were grouped into two classes, A/G and C/U. The paired cryoEM density and label volumes were partitioned into 92,014 subvolumes of 64³ Å³, which were subsequently divided into training, validation, and test sets.

### Training the U-Net models

Three independent 19-layer 3D U-Net models were trained to predict complementary structural and sequence-informative features from cryoEM density maps. The three models were designed to predict (1) structure classes, (2) key-atom positions, and (3) reduced residue or nucleotide types, respectively. Each model received a 64³ Å³ cryoEM density sub-volume as input and produces voxel-wise probability maps for its corresponding prediction classes (Supplementary Fig. 2).

All three models were trained using the Adam optimizer [36] with an initial learning rate of 1 × 10⁻⁴ and a batch size of 16. To improve convergence during later stages of training, we used a ReduceLROnPlateau learning-rate scheduler, which reduced the learning rate by a factor of 0.5 if the validation loss did not improve for five consecutive epochs. A weighted cross-entropy loss was used to address class imbalance, with class weights set according to the inverse frequency of each class. Training was performed on an NVIDIA GeForce RTX 4090 GPU with 24 GB of memory. Early stopping with a patience of 15 epochs was applied to reduce overfitting.

Model performance was monitored on the validation set throughout training, and the best-performing checkpoint for each model was retained for downstream evaluation. The selected checkpoints corresponded to epochs 76, 11, and 64 for the structure-class, key-atom, and reduced residue/nucleotide-type models, respectively.

### Evaluation of U-Net prediction performance

The outputs of the U-Net models are voxel-wise probability maps, with each voxel assigned a probability distribution over the corresponding prediction classes. For evaluation, a voxel was assigned to a non-background class when the corresponding predicted probability exceeded 0.8; otherwise, it was assigned to the background class. Ground-truth voxel labels were generated from the corresponding deposited atomic models using the labeling procedure described above.

Prediction performance was evaluated using voxel-wise accuracy, precision, recall, and F1 score. For a given class, true positives (TP), false positives (FP), false negatives (FN), and true negatives (TN) were defined by comparison with the corresponding ground-truth labels. The metrics were calculated as:

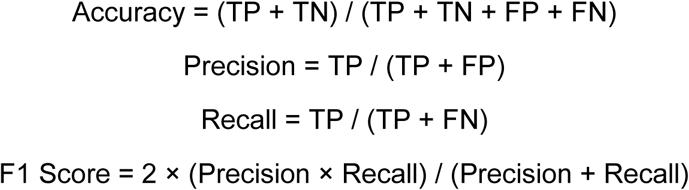

These metrics were calculated for the structure-class, key-atom, and reduced residue/nucleotide-type prediction models and were used to quantify voxel-level prediction performance.

### Backbone tracing and query generation

For backbone reconstruction, key-atom probability voxels predicted by the key-atom U-Net were first thresholded and grouped using mean-shift clustering [37]. The centroid of each cluster was treated as a predicted key-atom position. For protein backbone reconstruction, Cα positions served as key-atoms, whereas for RNA backbone reconstruction, both predicted sugar and phosphate positions served as key-atoms. These predicted key-atom positions were then used as graph nodes, and a radius-based nearest-neighbor graph was constructed by connecting nodes within a predefined distance threshold, producing an initially overconnected graph that contained geometrically plausible backbone connections.

The initial graph was subsequently refined using a U-Net-predicted secondary-structure field as a soft structural scaffold. For protein reconstruction, the predicted helix field was used, while for RNA reconstruction, the predicted RNA secondary-structure field was used. Secondary-structure probability voxels were first thresholded to retain high-confidence predictions. For each candidate edge, a scaffold-support score was calculated from the predicted secondary-structure signal within a cylindrical region surrounding the edge and normalized by edge length. The score is

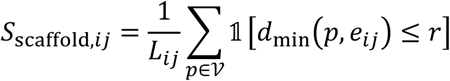

where *i* and *j* are connected nodes in the graph, *e_ij_* is the edge between nodes *i* and *j*, *L_ij_* is the Euclidean length of edge *e_ij_*, V is the set of coordinates of voxels belonging to the thresholded secondary-structure region, *p* is is the Cartesian coordinate of a voxel in V, *d*_min_(*p*, *e_ij_*) is the minimum Euclidean distance from voxel coordinate *p* to edge *e_ij_*, *r* is the radius of the cylindrical region surrounding the edge, and 1[⋅] is the indicator function which equals 1 when the condition is true and 0 otherwise.

Candidate edges were ranked by this score and greedily retained subject to a maximum node degree of two, such that each predicted Cα position could be connected to at most two neighboring nodes. This procedure preferentially retained backbone connections supported by both geometric proximity and the predicted secondary-structure field.

Although edge pruning limits each node to a maximum degree of two, it can leave small cycles in the graph. These cycles introduce closed loops that prevent the graph from representing continuous, non-branching backbone traces. They were therefore addressed through an iterative repair procedure. Edges that produced cycles were replaced, when possible, with alternative candidate connections from the initial overconnected graph. Candidate repairs were evaluated using a combination of scaffold support and graph continuity. Scaffold support favors connections consistent with the predicted secondary-structure regions, while graph continuity favors connections that preserve larger local graph neighborhoods.

Graph continuity was scored using this formula:

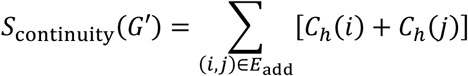

Here *G*′ is the candidate repaired graph being evaluated. *E*_add_ is the set of edges newly added during graph repair. (*i*, *j*) represents a newly added edge connecting nodes *i* and *j*. *C*_ℎ_(*i*) and *C*_ℎ_(*j*) are the numbers of distinct nodes reachable from nodes *i* and *j*, respectively, within ℎ graph hops, excluding the node itself. ℎ is the maximum number of graph hops considered when evaluating local connectivity, which is currently set at 8. *C*_ℎ_(*v*) for some arbitrary node *v* is defined mathematically as the following:

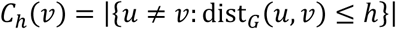

where *u* is another node in the graph, dist*_G_*(*u*, *v*) is the shortest-path distance between nodes *u* and *v* measured in graph hops, and ℎ is as defined above.

The scaffold and continuity scores were combined in the following manner:

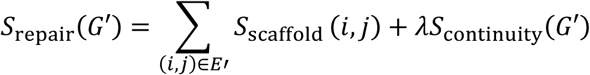

where *λ* is a weight, currently set at 0.25.

The refined graph was separated into connected components, and components below a minimum size threshold were discarded. Missing or unrecovered Cα positions naturally produced gaps in the reconstructed backbone, resulting in multiple disconnected backbone segments rather than a single continuous trace. Long connected components could optionally be divided into shorter segments to generate sequence queries of a desired maximum length.

Reduced residue types were then assigned to the graph nodes. For each node, reduced residue-type probabilities predicted by the U-Net were sampled within a local radius surrounding its position, and the residue class with the highest local probability was assigned. Each connected backbone component was then converted into a reduced-alphabet sequence and used as a query for downstream sequence-based candidate ranking. The resulting queries were exported as FASTA records.

### Sequence search module

For protein candidate ranking, map-derived query sequences and candidate protein sequences are recoded into the same six-class reduced amino acid alphabet: GASCVTI→G, P→P, LDNEQM→Z, KR→M, HFY→Y, and W→W. Thus, the G, P, L, K, Y, and W class labels exported by query generation become G, P, Z, M, Y, and W for searching. The mapper preserves X. Reduced candidate sequences are formatted as a local protein database using the software’s bundled NCBI BLAST+ 2.17.0 makeblastdb. Each candidate is assigned a unique internal database identifier that is mapped back to its original FASTA identifier in the results.

Duplicate candidate FASTA identifiers are rejected. Each query is searched together with its sequence reversal using gapped blastp-short [23,38], with the PAM30 substitution matrix, word size 2, E-value cutoff 1.0, gap-opening penalty 15, and gap-extension penalty 3. Composition-based statistics are disabled.

Protein alignments are retained if they cover at least 60% of the query length and the full candidate protein contains 1-100,000 residues. Gaps are allowed under the specified opening and extension penalties. For each query and candidate protein, the retained alignment with the lowest E-value is selected from searches using the query in both forward and reversed order. Different queries may align to separate or overlapping regions of the same candidate protein.

For protein sequence candidate j of length L_j_, let E_ij_ be the minimum retained E-value for query i, capped at 1.0, with E_ij_ = 1.0 when there is no retained hit. Among candidates with at least one retained alignment, the composite statistic is

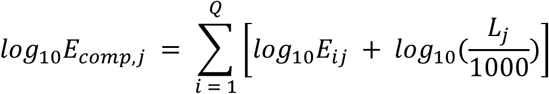

where Q is the number of original query fragments. Forward and reversed copies contribute a single term per fragment. The length factor is applied once for every query, including queries without retained hits. Calculations and sorting use logarithmic values to avoid numerical underflow or overflow in the product. For a positive BLAST-reported E-value, its base-10 logarithm is used directly. If BLAST reports an E-value of zero, the logarithm is recovered from the alignment bit score and the explicitly supplied search space as log10(E_ij_) = log10(Dq_i_) – B_ij_ log10(2). Here, D is the total number of residues in the candidate database, q_i_ is the length of query i, and B_ij_ is the BLAST bit score of the selected query-candidate alignment, where a larger bit score indicates a stronger match. Candidates are ordered by ascending composite log E-value. The primary protein score written on the software’s display sequence-ranking table is −log10(E_comp_,j), so larger reported primary scores correspond to stronger support.

For RNA candidate ranking, map-derived query fragments and candidate RNA sequences were recoded into the same two-class reduced nucleotide alphabet, with A and G mapped to A and C, T, and U mapped to U. Each query was compared with every candidate using ungapped sliding-window matching in both forward and reversed query orientations. For each query-candidate pair, the maximum fraction of matching reduced nucleotide symbols across all possible alignments and both query orientations was retained as the local sequence identity.

For each query, candidates are then ranked by local sequence identity. These ranks are divided by the number of candidate records to produce percentile scores, with higher values indicating better matches relative to the pool. Tied identities receive the average of their occupied rank positions before division. Each candidate’s final score is its lowest percentile across all queries. This is termed maximin scoring because it ranks candidates according to their weakest relative match. An example calculation can be found in Supplementary Table 4.

### RNA candidate-pool construction

For the Ca-raiA proof-of-concept analysis, the RNA candidate pool was constructed from the complete *Clostridium acetobutylicum* ATCC 824 genome assembly GCF_000008765.1, including chromosome NC_003030.1 and plasmid NC_001988.2 [32], together with the corresponding genome annotation. Because the target was known to be a non-coding RNA, annotated coding sequence (CDS) regions were excluded from the genomic sequence.

Contiguous non-CDS intervals of at least 50 nucleotides were retained. Each interval was represented by both its forward sequence and reverse complement, and thymine was converted to uracil. This procedure generated 4,632 candidate sequence records corresponding to 2,316 genomic intervals.

For this proof-of-concept analysis, the candidate pool was further constrained using the known 204-nucleotide reference target length. Candidate records between 102 and 306 nucleotides, corresponding to ±50% of the reference length, were retained. The resulting search pool contained 2,538 sequence records representing 1,269 genomic intervals.

### Structural retrieval

Structural retrieval uses the full candidate sequences in the saved sequence-ranking table. By default, the highest-ranked protein sequence or the top 50% of ranked RNA records are selected. The selection can be configured as an integer number of records or a decimal fraction of the ranked list. At the start of protein structural retrieval, the pipeline downloads RCSB PDB pdb_seqres.txt.gz. Records annotated mol:protein are retained, and identical protein sequences are deduplicated while preserving all associated PDB and chain identifiers. Each selected full protein sequence is aligned against every unique database sequence using the parasail striped Smith-Waterman implementation [39,40], BLOSUM62 [41], a gap-opening penalty of 11, and a gap-extension penalty of 1. All unique sequences tied for the maximum alignment score are retained, and their associated PDB identifiers are deduplicated to form the experimental-model candidate list.

For each candidate PDB identifier, the downloader requests biological assembly 1 in mmCIF format. The complete downloaded coordinate file is fitted as one rigid object. The pipeline also obtains PDBe UniProt cross-references specifically for the chains associated with the tied highest-scoring sequences [27,42]. The resulting unique UniProt accessions are queried against the AlphaFold Database prediction API. All distinct model-file URLs returned for those accessions are considered. Available models are downloaded and fitted individually.

The RNA retrieval works by constructing a nucleotide BLAST database from pdb_seqres.txt.gz retaining records annotated mol:na whose descriptions contain RNA and exclude DNA. Each selected full RNA sequence is searched using the bundled BLAST+ 2.17.0 [23] with -task blastn, -evalue 1e-3, and -max_target_seqs 100. Reported hits are filtered to query coverage greater than or equal to 30%. Hits for each candidate are ordered by descending bit score, where higher values indicate stronger matches. The three top scoring hits that have not already been selected anywhere in the run proceeds to alignment.

### Fitting fields and scoring

For a rigid transformation T(x) = Rx + t, the refinement and final ranking score is

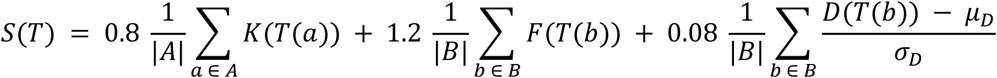

Here, T(x) = Rx + t describes how the model is placed in the map. R rotates an atom’s original position x, and t shifts it. The same transformation is applied to every atom, preserving the model’s shape. The sets A and B contain the atom positions used for scoring, and |A| and |B| are each set’s respective atom counts. For protein models, set A contains all Cα positions, whereas B contains all backbone N, Cα, C and O positions. The key field K is the predicted Cα probability map, and the scaffold field F is the max of the predicted helix, sheet and coil probabilities at each voxel. For RNA, set A contains all P positions, whereas B contains all N, CA, C, O or P positions. The key field is the phosphate probability map, and the scaffold field is the RNA structural probability map. Thus, K(T(a)) is the predicted probability at atom a’s transformed position, while F(T(b)) and D(T(b)) are the scaffold probability and experimental density at atom b’s transformed position. Each map is sampled at the transformed positions using trilinear interpolation, which takes a distance-weighted average of the eight surrounding voxels. Positions outside the grid use the nearest boundary value. Experimental density is standardized with z-scores by subtracting the whole-map mean μ_D_ and dividing by the whole-map population standard deviation σ_D_. The formula averages each signal over its atom set and combines the averages with manual tuned weights of 0.8, 1.2 and 0.08. The higher the final score S, the better agreement with these signals. The initial coarse composite score ranking uses a density weight of 0.04 instead of 0.08.

### Initial poses and rigid-body refinement

Initial model placements are generated by aligning the overall shape of the model’s atom positions with high-probability regions of predicted maps K and F. Both maps are evaluated at both the 99.95th and 99.50th percentiles, corresponding approximately to their highest-valued 0.05% and 0.50% of voxels, respectively. At each threshold, the selection is capped at the 12,000 highest-valued voxels for the key-atom map and 16,000 for the scaffold map. This strategy, along with its numbers and percentiles, was chosen because it quickly generated a vast number of initial poses that were roughly aligned with the overall density. The center and three principal axes of each selected voxel cloud are calculated, with higher-probability voxels receiving greater weight. Corresponding centers and principal axes are calculated from the model’s key-atom positions for comparison with the key-atom map, or backbone scoring positions for comparison with the scaffold map, giving each atom equal weight. These two maps and two thresholds produce four starting-pose configurations at the two different density thresholds. For each configuration, the model is rotated through 24 axis alignments that preserve its handedness and translated to align the model and voxel-cloud centers. The resulting 96 candidate placements are evaluated using the coarse composite score S. Only the highest-scoring placement proceeds to refinement.

The selected pose first undergoes eight iterations of deterministic point-cloud matching. Target points are taken from the 99.95th percentile of K and F, capped at 600 and 900 points, respectively. Each target point is paired with its nearest transformed model atom in the corresponding atom set. The closest 70% of matches from each field are retained, rounding upward, while keeping at least three key-field matches and six scaffold-field matches whenever available. The combined correspondences define a Kabsch least-squares rigid transformation [43]. The resulting pose is then optimized against S using bounded six-degree-of-freedom Powell optimization [44], with rotation about the centroid of B. Each rotation-vector component is bounded by ±45°, and each translation component by ±0.35L, where L is the longest physical extent of the experimental map. Powell optimization used parameter-search and relative score-improvement tolerances of 10^-6^, with a maximum of 96 iterations. The better of the starting and optimized poses under S is retained as the final model alignment. Of all candidate models aligned, the highest scoring AlphaFold and experimental model are returned.

### Synthetic-density controls

After establishing the protein identity as PfA-M17, we performed two synthetic-density controls to evaluate AlphaFold Database model retrieval and structural fitting (Fig. 5a). We generated maps at 2.7 Å resolution using the “molmap” command in UCSF ChimeraX 1.9 [24], retaining all other default settings. The first map used the 3,960 protein atoms of the 519 resolved polymer residues in PDB 4ZX8 chain A [29]. The second used all 4,789 atoms of AlphaFold Database model A0A024V0B1 [21]. Both maps retained their source models’ original coordinate frames and were saved in MRC format. We then processed each map with cryoID2 using the same malaria candidate sequence pool.

### Quantification and statistical analysis

Voxel-level feature prediction performance was summarized using accuracy, precision, recall, and F1 score. Cα-clustering precision, recall, and F1 score, as well as average protein-query sequence identity, were evaluated as functions of cryoEM map resolution. The relationship between average query identity and map resolution was summarized using ordinary least-squares linear regression and the corresponding coefficient of determination (R²). No inferential significance test was applied.

Structural retrieval and fitting were evaluated using 52 independent cryoEM map-model cases in each of two benchmark modes. Deposited-model and fitted-model Q-scores [28] were paired by case. For benchmarking, starting poses were randomized by applying a random rotation followed by an in-bound translation with a 4 Å margin. Production fitting of experimental cases did not include this starting-pose randomization.

### Benchmark dataset

CryoID2 was evaluated using a curated dataset of 52 published protein cryoEM cases obtained from the Electron Microscopy Data Bank (EMDB) [45]. The dataset included proteins from diverse functional classes, including ion channels, transporters, enzymes, soluble protein assemblies, bacterial proteins, and molecular chaperones, spanning a range of oligomeric states and cryoEM map resolutions.

For each case, the benchmark dataset included the target protein sequence, the corresponding deposited cryoEM density map, a deposited PDB atomic model used as the ground truth structural reference, the reported map resolution and recommended contour level, and precomputed prediction maps defined on the same voxel grid as the experimental density. In this case, these prediction maps were generated by the cryoID2 feature-prediction module and were treated as fixed inputs for the structural retrieval and alignment benchmark.

## Notes

### Competing Interest Statement

The authors have declared no competing interest.

