## Supplementary material for "CryoID2: deep learning-enhanced identification of protein and RNA from cryoEM maps": ALL Supplementary info

### Supplementary Data for CryoID2: deep learning-enhanced identification of protein and RNA from cryoEM maps

### Supplementary Table 1

| Gene | GT PDB | Fetch selected | GT Q-score | Fetch Q-score |
|---|---|---|---|---|
| vipp1 | 6ZW4 | PDB 6ZW4 | 0.3941 | 0.3941 |
| xptA2 | 9MLI | PDB 9MLI | 0.6697 | 0.6672 |
| VP1 | 9LGJ | PDB 9LGJ | 0.7708 | 0.7708 |
| clpL | 6LSY | PDB 6LSY | 0.2348 | 0.2349 |
| POLQ | 9BH7 | PDB 9BH7 | 0.5806 | 0.5805 |
| MUF8.1 | 6LBA | PDB 6LBA | 0.5377 | 0.5377 |
| clhm-1 | 6LOM | PDB 6LOM | 0.4594 | 0.4594 |
| cdtB | 6OKR | PDB 6OKR | 0.5831 | 0.5817 |
| Trpv2 | 5HI9 | PDB 5HI9 | 0.3223 | 0.3223 |
| GSDMB | 8ET2 | PDB 8ET1 | 0.1754 | 0.1908 |
| OGA | 9NE2 | PDB 9NE2 | 0.5084 | 0.4977 |
| Panx1 | 9HEV | PDB 9HEV | 0.2947 | 0.2943 |
| ribH | 9G3J | PDB 9G3J | 0.5909 | 0.5908 |
| LRRC8A | 6DJB | PDB 6DJB | 0.3892 | 0.1746 |
| RYR1 | 8X48 | PDB 8SEY | 0.4671 | 0.4598 |
| gdh | 9HV0 | PDB 9HV0 | 0.6999 | 0.6999 |
| vat | 5G4F | PDB 5G4F | 0.2419 | 0.242 |
| ALB | 8VAF | PDB 8VAF | 0.6122 | 0.6121 |
| TRPV6 | 6D7S | PDB 6D7S | 0.4786 | 0.4786 |
| clpC | 6EM9 | PDB 6EM9 | 0.2576 | 0.2577 |
| TTR | 9ZBV | PDB 9ZBV | 0.6583 | 0.6435 |
| mthK | 6UXA | PDB 6UX4 | 0.4151 | 0.3517 |
| APOB | 9E9R | PDB 9BD1 | 0.247 | 0.022 |
| im30 | 9I00 | PDB 8AKZ | 0.3074 | 0.198 |
| Slc26a4 | 7WL7 | PDB 7WL7 | 0.4558 | 0.4559 |
| GJB2 | 6UVS | PDB 6UVR | 0.5306 | 0.4757 |
| mscS | 6UZH | PDB 6UZH | 0.6599 | 0.6593 |
| ALMT1 | 7VQ3 | PDB 7VQ3 | 0.6749 | 0.675 |
| corA | 3JCH | PDB 3JCH | 0.4227 | 0.4226 |
| KCNMA1 | 6ND0 | PDB 6ND0 | 0.5485 | 0.5485 |
| mexB | 6T7S | PDB 6T7S | 0.4302 | 0.3582 |
| chdC | 8QWC | PDB 6XUB | 0.7829 | 0.6511 |
| AUX1 | 9UTE | PDB 9UTE | 0.7 | 0.6998 |
| NF1 | 7PGP | PDB 7PGP | 0.6858 | 0.6855 |
| ANO10 | 6R7Z | PDB 6R7Z | 0.3949 | 0.3948 |
| TRPV3 | 8GKG | PDB 8GKG | 0.4294 | 0.0177 |
| Grik1 | 8GPR | PDB 7YSV | 0.1964 | 0.1914 |
| flaB2 | 9X7K | PDB 9X7M | 0.3628 | 0.2115 |
| ABCB6 | 7D7R | PDB 7D7R | 0.4049 | 0.4049 |
| DERL1 | 7CZB | PDB 7CZB | 0.5327 | 0.5329 |

| | | | | |
|---|---|---|---|---|
| CRYAB | 3J07 | PDB 2KLR | 0.0322 | 0.1192 |
| ABCC8 | 6PZC | PDB 6PZC | 0.4469 | 0.4469 |
| ABCD1 | 7RRA | PDB 7RRA | 0.3186 | 0.3184 |
| SPOP | 9OUT | PDB 9OUT | 0.2854 | 0.2853 |
| CLCN1 | 6QVD | PDB 6QV6 | 0.3331 | 0.3374 |
| Kcnj6 | 6XIS | PDB 6XIS | 0.4719 | 0.4719 |
| SLC2A9 | 9CAX | PDB 9CAX | 0.62 | 0.62 |
| Slc26a9 | 6RTF | PDB 6RTF | 0.3313 | 0.3312 |
| TOMM20 | 7VC9 | PDB 7VC9 | 0.127 | 0.1271 |
| fliC | 7SN7 | PDB 8CVI | 0.4665 | 0.039 |
| Ano1 | 5NL2 | PDB 5NL2 | 0.4325 | 0.4323 |
| Tmem63c | 8K0B | PDB 8K0B | 0.2474 | 0.247 |

### Supplementary Table 2

| Gene | GT Q-score | Aligned Q-score |
|---|---|---|
| vipp1 | 0.3941 | 0.3941 |
| xptA2 | 0.6697 | 0.6672 |
| VP1 | 0.7708 | 0.7708 |
| clpL | 0.2348 | 0.2349 |
| POLQ | 0.5806 | 0.5805 |
| MUF8.1 | 0.5377 | 0.5377 |
| clhm-1 | 0.4594 | 0.4594 |
| cdtB | 0.5831 | 0.5817 |
| Trpv2 | 0.3223 | 0.3223 |
| GSDMB | 0.1754 | 0.1747 |
| OGA | 0.5084 | 0.4977 |
| Panx1 | 0.2947 | 0.2943 |
| ribH | 0.5909 | 0.5908 |
| LRRC8A | 0.3892 | 0.1746 |
| RYR1 | 0.4671 | 0.4671 |
| gdh | 0.6999 | 0.6999 |
| vat | 0.2419 | 0.242 |
| ALB | 0.6122 | 0.6121 |
| TRPV6 | 0.4786 | 0.4786 |
| clpC | 0.2576 | 0.2577 |
| TTR | 0.6583 | 0.6435 |
| mthK | 0.4151 | 0.4152 |
| APOB | 0.247 | 0.0247 |
| im30 | 0.3074 | 0.307 |
| Slc26a4 | 0.4558 | 0.4559 |
| GJB2 | 0.5306 | 0.5309 |
| mscS | 0.6599 | 0.6593 |
| ALMT1 | 0.6749 | 0.675 |
| corA | 0.4227 | 0.4226 |
| KCNMA1 | 0.5485 | 0.5485 |
| mexB | 0.4302 | 0.3582 |
| chdC | 0.7829 | 0.1418 |
| AUX1 | 0.7 | 0.6998 |
| NF1 | 0.6858 | 0.6855 |
| ANO10 | 0.3949 | 0.3948 |
| TRPV3 | 0.4294 | 0.0177 |
| Grik1 | 0.1964 | 0.1964 |
| flaB2 | 0.3628 | 0.2188 |
| ABCB6 | 0.4049 | 0.4049 |
| DERL1 | 0.5327 | 0.5329 |

| | | |
|---|---|---|
| CRYAB | 0.0322 | 0.066 |
| ABCC8 | 0.4469 | 0.4469 |
| ABCD1 | 0.3186 | 0.3184 |
| SPOP | 0.2854 | 0.2853 |
| CLCN1 | 0.3331 | 0.3332 |
| Kcnj6 | 0.4719 | 0.4719 |
| SLC2A9 | 0.62 | 0.62 |
| Slc26a9 | 0.3313 | 0.3312 |
| TOMM20 | 0.127 | 0.1271 |
| fliC | 0.4665 | 0.4665 |
| Ano1 | 0.4325 | 0.4323 |
| Tmem63c | 0.2474 | 0.247 |

### Supplementary Table 3

Supplementary Table 3 is provided as a separate file because of its large size.

### Supplementary Table 4

| Candidate | Query 1 local identity | Query 1 percentile | Query 2 local identity | Query 2 percentile | Final score (lower percentile) |
|---|---|---|---|---|---|
| A | 90% | 1 | 50% | 0.25 | 0.25 |
| B | 80% | 0.75 | 80% | 1 | 0.75 |
| C | 70% | 0.5 | 70% | 0.75 | 0.5 |
| D | 60% | 0.25 | 60% | 0.5 | 0.25 |

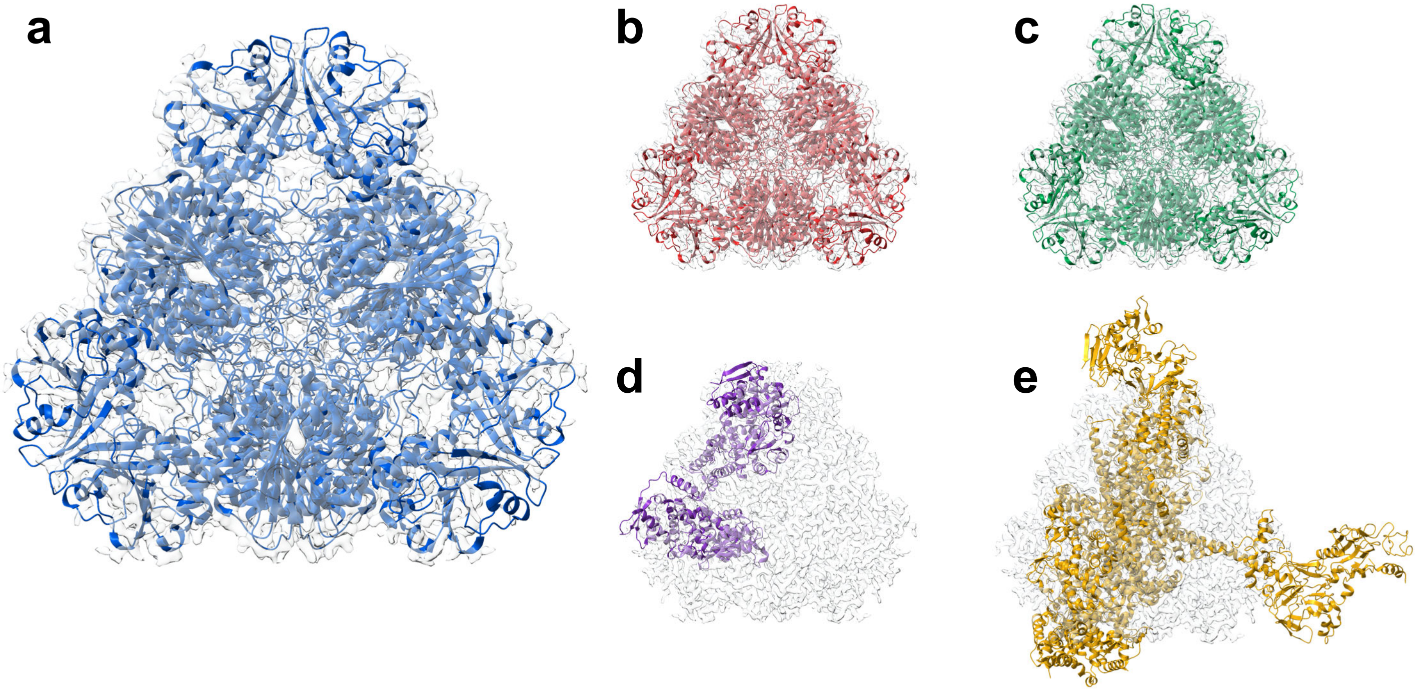


**Supplementary Figure 1. Ranked models fitted to the E1_D3 cryo-EM density map.** (a) First-ranked model for the first-ranked protein sequence (PDB: 4ZX8). (b) Second-ranked model for the first-ranked protein sequence (PDB: 4ZY1). (c) Third-ranked model for the first-ranked protein sequence (PDB: 4ZX9). (d) First-ranked model for the second-ranked protein sequence (PDB: 5LP8). (e) First-ranked model for the third-ranked protein sequence (PDB: 9YGJ).

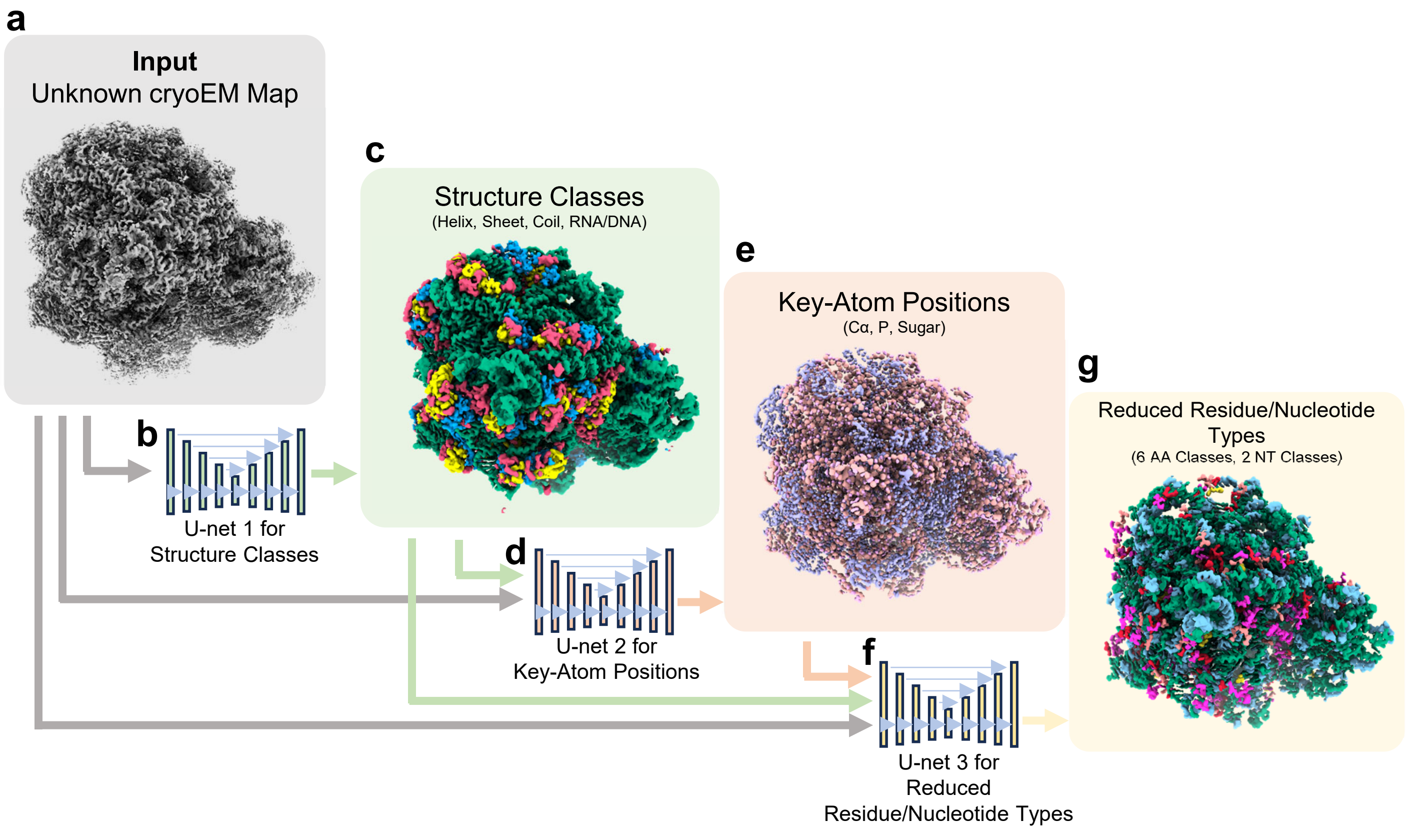


**Supplementary Figure 2. Architecture and outputs of the three cryoID2 U-Net models.** **(a)** An input cryoEM density map is processed sequentially by three 19-layer 3D U-Net models. **(b,c)** U-Net 1 predicts voxel-wise structure classes, including helix, sheet, coil, RNA/DNA, and background. **(d,e)** U-Net 2 predicts key-atom positions, including protein Cα atoms, RNA phosphate atoms, and RNA sugar-ring atoms. **(f,g)** U-Net 3 predicts reduced residue or nucleotide types. Protein residues are represented by six reduced amino-acid classes, whereas RNA nucleotides are represented by two reduced nucleotide classes. The outputs of the preceding prediction stages are incorporated into the downstream U-Net inputs as indicated by the arrows.
